# High conformational flexibility of phosphomannomutase 2: Implications for functioning mechanisms, stability and pharmacological chaperone design

**DOI:** 10.1101/2025.01.27.635082

**Authors:** Francisco del Caño-Ochoa, Marçal Vilar, Alicia Vilas, Rebeca Company, Belén Pérez, Santiago Ramón-Maiques

**Author notes:** Corresponding author: Santiago Ramón-Maiques, Instituto de Biomedicina de Valencia (IBV), CSIC, Jaime Roig, 11. Valencia, 46010, Spain., (ext. 431468).

## Abstract

Phosphomannomutase 2 (PMM2) is a critical enzyme in the N-glycosylation pathway, and its defect is the cause of the most common congenital disorder of glycosylation. Despite its biological relevance, the understanding of PMM2 is limited, as the catalytic mechanism and required conformational dynamics remain unknown. In this study, we investigated murine PMM2 (Pmm2) to elucidate its structural flexibility and functional insights. High-resolution crystal structures of Pmm2 in both apo and activator-bound forms provided a more detailed model of the protein, underscoring the role of three ionic cofactors that are essential for dimerization, catalysis, and stability. The Pmm2 structures also provided eight distinct conformations of the protein subunit, highlighting its dynamic nature. Structural comparisons among Pmm2, human PMM2 and other phosphomannomutases helped define the architecture of the enzyme as a dimer assembled by the rigid association of the cap domains, which provide a flat platform from which the core domains of each subunit protrude in a flexible manner. Molecular dynamics (MD) simulations of the human and murine PMM2s further emphasized the enzyme’s substantial conformational flexibility, revealing extensive core domain movements and suggesting potential inter-subunit communication within the dimer. This study refines the model of PMM2 function, demonstrating its dynamic role in substrate binding, intermediate reorientation, and product release. The observed flexibility provides new opportunities to target specific enzyme states, enabling the development of pharmacological chaperones.

## Introduction

Defects in the enzyme phosphomannomutase 2 (PMM2; E.C.: 5.4.2.8) impair protein N-glycosylation and cause the most common form of congenital disorder of glycosylation (CDG) (Jaeken, 2010; Jaeken and Péanne, 2017; Matthijs et al., 1997; Ng and Freeze, 2018; Van Schaftingen and Jaeken, 1995). PMM2-CDG (MIM# 212065) is a rare autosomal recessive disorder that affects the nervous system and other organs, often resulting in fatality during the first years of life, although there have been reports of patients with milder symptoms (Freeze, 2009; Giurgea et al., 2005; Grünewald, 2009). At present, there are no specific therapies available for PMM2-CDG patients beyond symptomatic treatments, and discovering a cure remains a major challenge (Gámez et al., 2020). More than 100 pathogenic variants have been documented for *PMM2* (Human Gene Mutation Database; http://www.hgmdl.cf.ac.uk) (Stenson et al., 2014). Most of these variants compromise protein stability, leading to decreased enzymatic activity caused by reduced lifetimes and lower cellular concentrations (Vega et al., 2011; Yuste-Checa et al., 2015). A potential therapeutic approach for PMM2-CDG involves using chemical compounds as pharmacological chaperones that stabilize PMM2 variants with folding defects (Gámez et al., 2018; Vilas et al., 2020). As a proof-of-concept, *in vitro* and cellular assays have shown that 1-(3-chlorophenyl)-3,3-bis(pyridin-2-yl)-urea, referred to as “compound VIII”, enhances the stability and/or increases the residual activity of certain PMM2 folding variants. Nevertheless, the precise mechanism responsible for this effect remains unknown (Segovia-Falquina et al., 2022; Yuste-Checa et al., 2017). Thus, a thorough structural characterization of PMM2 would be crucial for understanding the folding defects associated to PMM2-CDG variants and developing stabilizing treatments. To this end, we determined the crystal structure of human PMM2 (hPMM2) homodimer (Briso-Montiano et al., 2022, p.). However, key aspects of PMM2 remain unknown, particularly regarding the reaction mechanism and the associated conformational movements.

PMM2 catalyzes the conversion of mannose-6-phosphate (Man6P) into mannose-1-phosphate (Man1P), the precursor of the GDP-mannose that serves as a high-energy nucleotide sugar donor for various mannosyltransferases (Matthijs et al., 1997; Pirard et al., 1999). The reaction involves a two-step phosphoryl transfer with the reorientation of a reaction intermediate in the process (Silvaggi et al., 2006) (Figure 1). Before starting the reaction, the enzyme is activated by transferring a phosphate group from α-glucose 1,6-bisphosphate (Glc-1,6-P_2_) to a specific aspartate residue within the active site (Collet et al., 1998). Subsequently, the activated enzyme transfers the phosphate group to the substrate Man6P, resulting in mannose-1,6-bisphosphate (Man-1,6-P_2_), which is an intermediate in the reaction. This intermediate may either be reoriented within the active site or released and rebound in the opposite orientation. Next, the aspartate attacks the reoriented Man-1,6-P_2_, producing Man1P, and the Asp remains phosphorylated for the subsequent reaction cycle. A series of conformational changes are thought to be essential for the sequential binding and release of the activator, substrate, and product, for enabling the reorientation of the reaction intermediate, and for preventing the incorporation of water molecules that could potentially act as phosphate acceptors. Evidence supporting these movements was gathered through the structural analysis of a related phosphomannomutase from *Leishmania mexicana* (leisPMM), which was crystallized in a combination of dimers with subunits in open and closed conformations (Kedzierski et al., 2006). Similar protein motions have been suggested for hPMM2 based on limited proteolysis, chemical modification assays, and molecular dynamics (MD) simulations of the protein subunit (Andreotti et al., 2014; Monticelli et al., 2019). However, the crystal structure of hPMM2 showed the protein exclusively in open conformations, even with Glc-1,6-P_2_ bound to the active site (Briso-Montiano et al., 2022, p.). Similarly, the crystal structures of human PMM1 (hPMM1), an isoenzyme that shares 64% sequence identity with hPMM2 and catalyzes the same reaction (though it does not compensate for hPMM2 deficiency) (Cromphout et al., 2006, 2005), consistently showed open conformations, whether Man1P or inosine monophosphate (IMP) was bound (Ji et al., 2018; Silvaggi et al., 2006). Consequently, the currently available structures offer limited insights into the enzyme’s flexibility, highlighting the need for additional structural studies to unravel the conformational changes crucial for its function.

**Figure 1.**
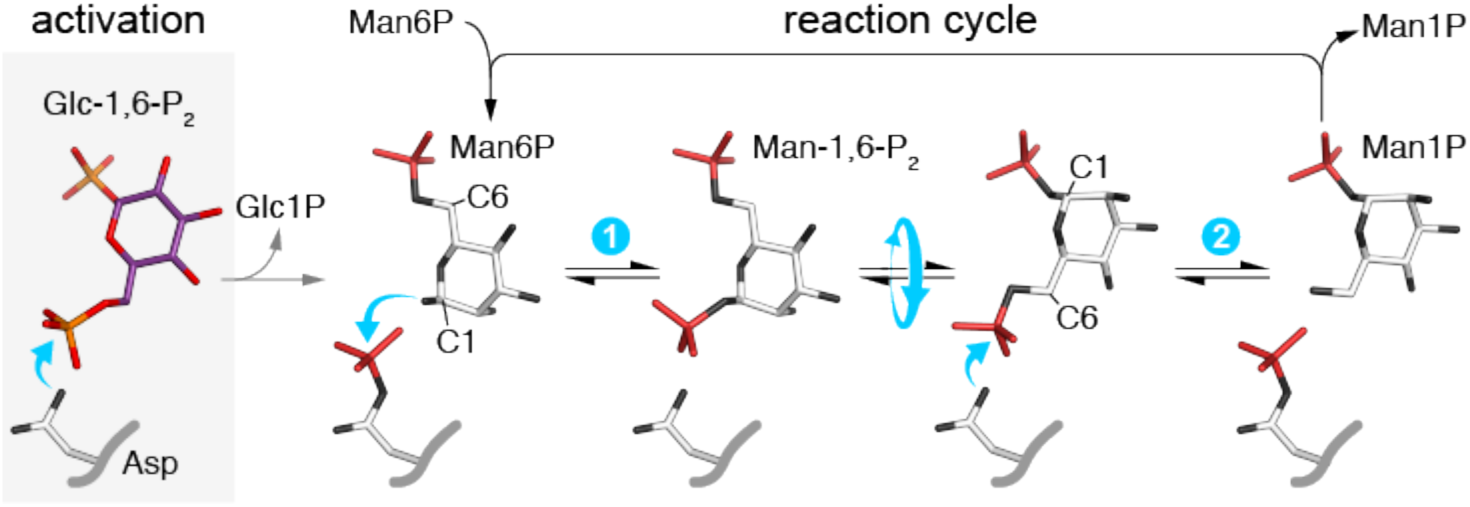
Reaction catalyzed by PMM2. The activation step is highlighted in grey background. The two phosphoryl transfer reactions are numbered. The curved cyan arrows represent nucleophilic attacks, and the round arrow indicates the reorientation of the reaction intermediate.

In this study, we present the crystal structure of mouse PMM2 (Pmm2) in its free form and bound to the activator Glc-1,6-P_2_. The structures, obtained from two different crystal forms, provide a higher-resolution view of the enzyme, showcasing eight distinct conformations and showing how essential ion co-factors coordinate for dimerization, catalysis, and protein stability. The comparison of the structures of mouse and human PMM2 and molecular dynamics (MD) simulations reveal a remarkable level of conformational flexibility. These findings raise questions about possible communication between subunits within the dimer and the catalytic mechanism and suggest potential conformational targets for pharmacochaperone design.

## RESULTS

### Production and crystal structure of mouse Pmm2

We expressed and purified the recombinant mouse Pmm2 (UniProt Q9Z2M7) using the same strategy previously utilized for hPMM2 (Briso-Montiano et al., 2022). The purified Pmm2 appeared as a single band in SDS-PAGE, with the expected molecular weight of 27.8 kDa (Figure 2A). Size-exclusion chromatography (SEC) analysis confirmed the formation of a stable homodimer in solution, with an estimated molecular weight of 51 kDa (Figure 2B,C). As previously reported (Segovia-Falquina et al., 2022), the SEC elution of Pmm2 was delayed compared to the hPMM2 dimer, with an estimated molecular weight of 61 kDa (Figure 2B top). Both proteins share 91% sequence identity (Supplementary Figure 1), and the four additional residues at the N-terminus of hPMM2 could not account for the apparent 10 kDa difference between dimers. Thus, the apparent smaller size of Pmm2 was attributed to a more compact conformation impacting SEC migration. Since hPMM2 was crystallized in an open conformation (Briso-Montiano et al., 2022), this result suggested that Pmm2 could be a good candidate for crystallizing the enzyme in a closed state, which led to the structural studies reported below. However, as explained later, structural studies suggested that Mg^2+^ binding was integral to protein folding, and the repetition of SEC assays with MgCl_2_ in the running buffer showed that both proteins eluted at the same position (Figure 2B bottom), confirming the key role of Mg^2+^ in protein structure.

**Figure 2.**
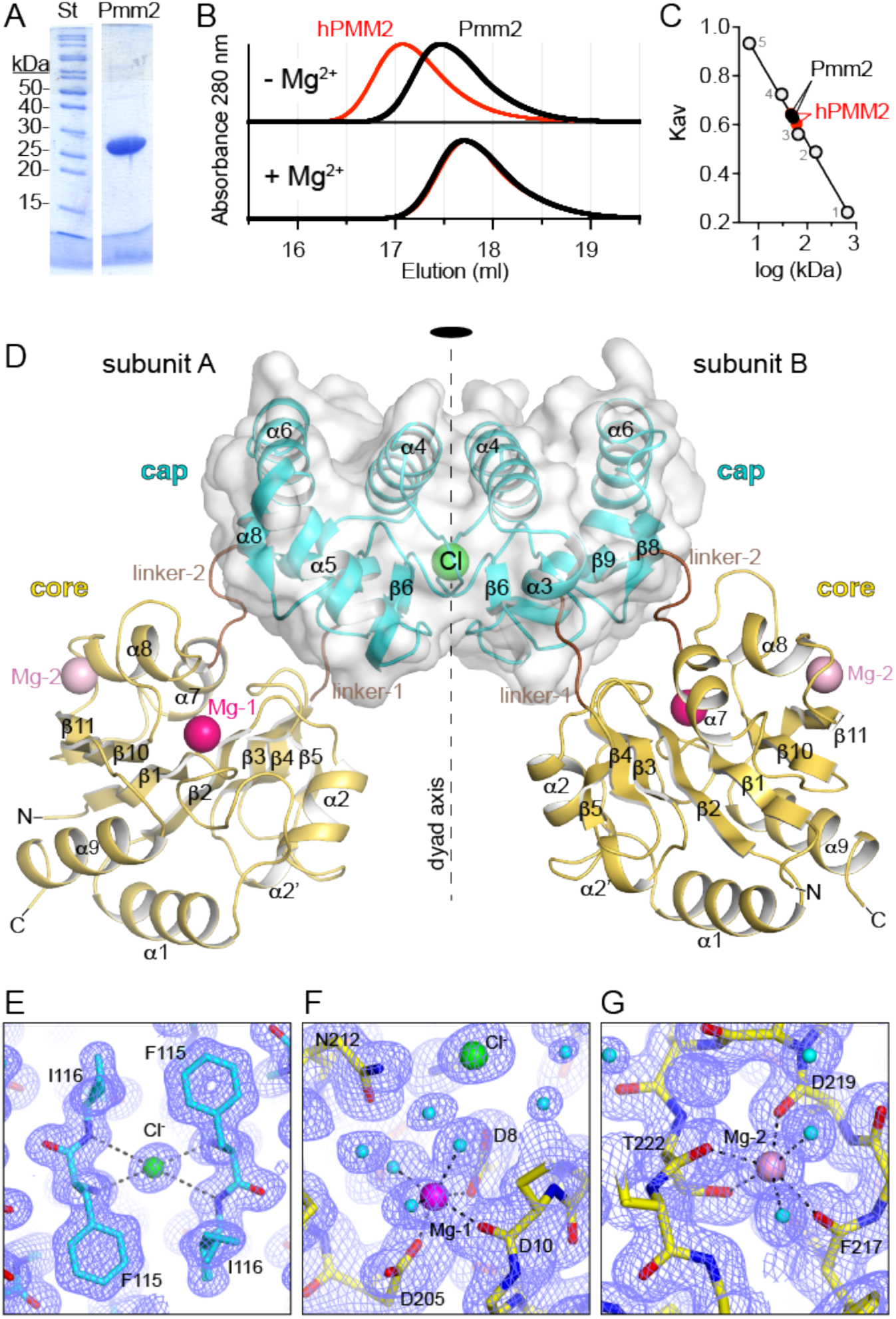
Crystal structure of mouse PMM2. **A**) Purified mouse Pmm2 on SDS-PAGE. St, molecular weight standards. **B**) Size exclusion chromatography elution profile of purified mouse (black trace) and human (red) PMM2. **C)** Column calibration with protein standards (1, ribonuclease A, 13.7 kDa; 2, carbonic anhydrase, 29 kDa; 3, conalbumin, 75 kDa; 4, aldolase, 158 kDa; and 5, ferritin, 440 kDa). **D**) Crystal structure of Pmm2 homodimer with the core and cap domains colored yellow and cyan, respectively, and the linker regions colored brown. The cap-cap dimer is shown in a semi-transparent surface representation to highlight its rigidity, and the embedded chlorine anion is represented as a green sphere. The Mg^2+^ ions in the core domains, Mg-1 and Mg-2, are shown respectively as magenta and pink spheres. **E–F)** Detail of the 2F_obs_-F_calc_ electron density maps contoured at 1.0 σ showing the coordination of the ionic cofactors. Dashed lines indicate the interactions of the ions with the protein.

We successfully grew two different types of Pmm2 crystals in conditions containing either PEG 8000 (referred to as type 1 crystal) or PEG 3350 (type 2 crystal) as the precipitant (Table 1). Type 1 crystals belonged to space group P2_1_2_1_2_1_, diffracted X-rays to 2.43 Å resolution, and contained six protein subunits (named A_1_–F_1_) arranged in three dimers (A_1_/B_1_, C_1_/D_1_, E_1_/F_1_) within the asymmetric unit. Type 2 crystals belonged to space group P6_5_22, diffracted X-rays to 1.49 Å resolution, and contained a dimer (A_2_/B_2_) within the asymmetric unit. Crystallographic phases were obtained by molecular replacement using hPMM2 as the search model. The electron density was well-defined and allowed to trace the entire polypeptide chains and various ions already seen in hPMM2, including a Cl^-^ anion in the dimerization interface, along with two Mg^2+^ cations, one at the active site (Mg-1), and the other (Mg-2) proposed to serve a stabilizing function (Briso-Montiano et al., 2022) (Figure 2D–G and Supplementary Figure S2A). An extra electron density indicated the binding of an additional Cl^-^ anion in the active site adjacent to Mg-1 (Figure 2F).

**Table 1.** Data collection and refinement statistics.

|  | **APO (PEG 3350)** | **APO (PEG 8000)** | **G16bP (PEG 8000)** |
| --- | --- | --- | --- |
| PDB ID | 9HN3 | 9HB | 9I55 |
| **Data Collection** |  |  |  |
| Beamline | BL13-XALOC (ALBA) | BL13-XALOC (ALBA) | BL13-XALOC (ALBA) |
| Wavelength (Å) | 0.97926 | 0.97926 | 0.97926 |
| Space group | P6<sub>5</sub>22 | P2<sub>1</sub>2<sub>1</sub>2<sub>1</sub> | P2<sub>1</sub>2<sub>1</sub>2<sub>1</sub> |
| Cell dimensions<br>*a*, *b*, *c* (Å) | 71.8, 71.8, 355.9 | 80.7, 98.6, 212.8 | 81.3, 99.1, 213.5 |
| Resolution (Å) | 177.95–1.49 (1.51–1.49)<sup>a,b</sup> | 106.43–2.43 (2.48–2.43) | 213.47–2.75 (2.85–2.75) |
| Total reflections | 1,721,710 (83,956) | 424,797 (19,107) | 298,159 (29,278) |
| Unique reflections | 89,547 (4,298) | 64,543 (3,193) | 45,273 (4,408) |
| R<sub>pim</sub> | 0.019 (0.627) | 0.046 (0.873) | 0.097 (0.883) |
| Mean I/σI | 16.7 (1.3) | 10.6 (0.9) | 6.6 (1.0) |
| Completeness (%) | 99.3 (98.2) | 99.9 (99.9) | 98.0 (100) |
| Multiplicity | 19.2 (19.5) | 6.6 (6.0) | 6.6 (6.6) |
| CC<sub>1/2</sub> | 0.999 (0.591) | 0.999 (0.614) | 0.993 (0.558) |
| **Refinement** |  |  |  |
| Resolution (Å) | 62.19–1.49 | 53.21–2.43 | 64.67–2.75 |
| No. of reflections used | 89,450 | 64,476 | 45,183 |
| R<sub>work</sub>/R<sub>free</sub> | 17.91/19.52 | 23.40/27.05 | 20.54/26.47 |
| Rmsd<br>Bond lengths (Å)<br>Bond angles (°) | 0.005<br>0.71 | 0.004<br>0.622 | 0.005<br>0.685 |
| No. of non-hydrogen atoms | 4,668 | 11,765 | 11,900 |
| Protein | 4,026 | 11,429 | 11,503 |
| Ligand | 38 | 19 | 151 |
| solvent | 618 | 317 | 294 |
| Ramachandran favored<br>allowed<br>outliers | 97.3<br>2.7<br>0 | 94.7<br>5.1<br>0.2 | 92.5<br>7<br>0.5 |
| Rotamer outliers (%) | 0.91 | 1.75 | 2.22 |
| Clashscore | 2.60 | 8.02 | 7.29 |
| Average B-factor<br>protein<br>ligands<br>solvent | 34.74<br>33.00<br>39.59<br>45.89 | 71.09<br>71.32<br>65.01<br>63.11 | 65.48<br>65.63<br>83.83<br>53.17 |

<sup>a</sup> Values in parenthesis correspond to the highest resolution shell.
<sup>b</sup> Criteria of CC<sub>1/2</sub> ≥ 0.5 was used in the determination of resolution cutoff.

### Mouse Pmm2 overall fold and conformations within the crystal

The overall fold of Pmm2 closely resembles that of hPMM2, but the higher resolution data, particularly from crystal type 2, enabled us to refine the protein model and better define the coordination of the ionic cofactors. The Pmm2 subunit is divided into a core domain (aa 1–79 and 185–242) holding the phosphoryl transfer site and a cap domain (aa 83–180) where the substrate binds in the first place, connected by two linker regions, linker-1 (aa 80–82) and linker-2 (aa 181–184) (Figure 2D and Supplementary Figures S1 and S2A). The globular core and cap domains exhibit a high degree of compactness and rigidity, as indicated by the low root-mean-square deviation (rmsd) values when superimposing the separated domains across the eight Pmm2 subunits observed in type 1 and type 2 crystals (core, rmsd=0.4–0.7 Å for 137 Cα atoms; cap, rmsd = 0.3–0.4 Å for 98 Cα’s) (Figure 3A).

**Figure 3.**
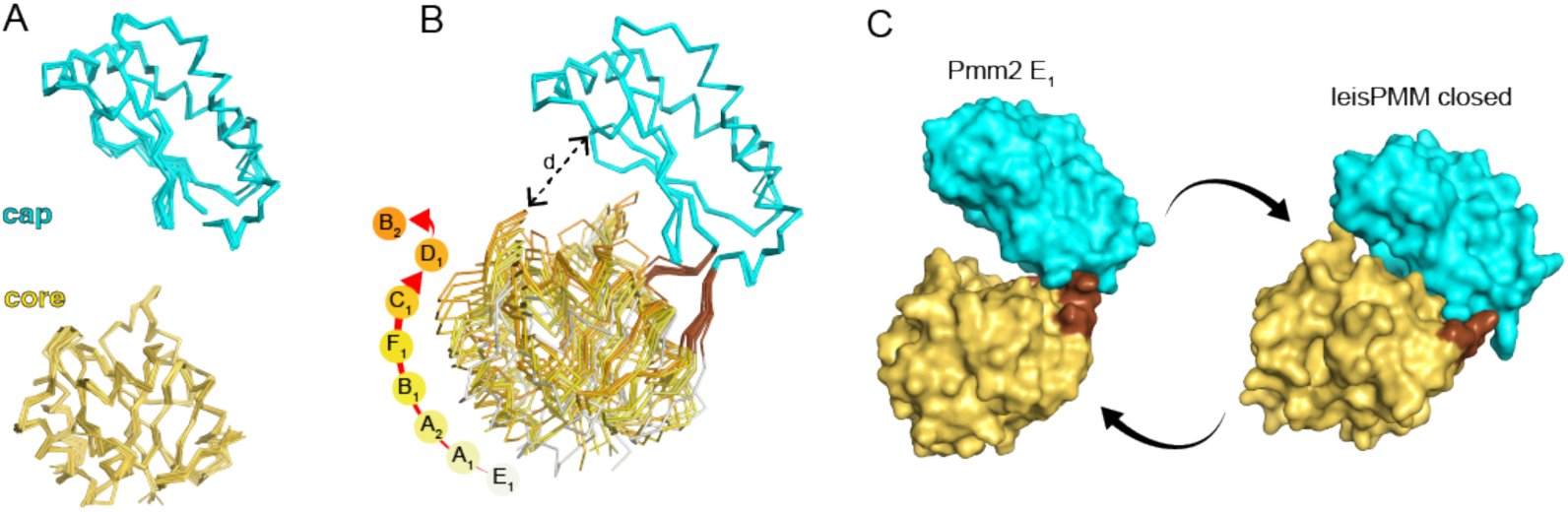
Pmm2 conformational movements. **A)** Ribbon representation of the independent superposition of the cap and core domains of the six Pmm2 subunits (A_1_–F_1_) in the asymmetric unit of the type 1 crystal, and the two Pmm2 subunits (A_2_ and B_2_) in the type 2 crystal. **B)** Superposition of the eight Pmm2 subunits through the cap domain, showing the different orientations of the core domains. The red arrow indicates the degree of closure of the different subunits that are colored from grey to orange. The dashed arrow indicates the distance between the Cα atoms of residues R21 in the core domain and Q176 in the cap domain. **C)** Surface representation of Pmm2 subunit E1 (left) and the closed conformation of leisPMM subunit (right). The arrows indicate the possible conformational movements between the open and closed conformations.

In the crystal structure, the dimerization of the protein exclusively relies on interactions between the cap domains, which form a 40 x 40 x 20 Å block with no significant variations among the four determined Pmm2 dimers (rmsd = 0.37–0.44 for 196 Cα’s) (Figure 3A and Supplementary Figure S2B). A virtually identical arrangement is found in the crystal structures of hPMM2, leisPMM, and hPMM1 (Supplementary Figure S2C). These findings support the view of the cap-cap dimer as a robust platform, represented in the figures with a semitransparent surface (Figure 2D and Supplementary Figure S2A), from which the core domains protrude.

The dimerization interface buries a Cl^-^ anion coordinated by the N atoms of residues F115 and I116 from both subunits (Figure 2D,E). The structures of leisPMM and hPMM2 also show the interposed Cl^-^ anion at the same position as observed in Pmm2, and although the structures of hPMM1 do not include the Cl^-^ anion in the final models (Ji et al., 2018; Silvaggi et al., 2006), analysis of the structural data strongly suggests the binding of the anion at the conserved position (Supplementary Figure S2C,D). Thus, the structures of all four Pmm2 dimers and the comparison with the homolog structures provide strong evidence supporting the Cl^-^ ion as a key cofactor in dimerization.

Importantly, there were significant differences in the relative orientations of the cap and core domains among the eight Pmm2 subunits observed in the two crystal forms (Figure 3B). These conformational differences can be explained by pure rotations of the core domains towards the cap domain, with a closure axis passing between the hinge residues in linker-1 and linker-2 (Figure 3B and Supplementary Figures S3 and S4). The D_1_ and E_1_ subunits show, respectively, the closest and most open conformations, with a 22° rotation between their core domains and a shortening of the distance to the cap domain from 22 Å to 14 Å (distance “d” measured between the Cα atoms of residues R17 and Q173) (Figure 3B and Supplementary Figure S4A). The comparison of all subunits shows roughly similar orientations for the rotation axes, suggesting a preferred trajectory for the subunit closure. However, subunit B_2_ deviates from this trajectory, suggesting a higher degree of freedom without a determined direction of the movement (Supplementary Figure S4).

### Conformational differences among PMMs crystal structures

To gain a better understanding of the enzyme’s flexibility, we compared the relative disposition of the core and cap domains within the structures of Pmm2, hPMM2, hPMM1, and leisPMM. Despite a moderate sequence identity of 45% (Supplementary Figure S5), these proteins exhibit high structural similarity in the cap and core domains (rmsd values < 1 Å) (Figure 3A), but differ in the relative orientations of the domains (Supplementary Figure S6A). As before, these differences can be described as rotations of the core domains around closure axes passing between the linker regions (Supplementary Figure S6B). Although this comparison does not suggest that the structures of the different enzymes correspond to conformations along a preferred closure path, it is worth noting that the superposition of leisPMM with Pmm2 shows a rotation axis with roughly similar orientation to that observed between the Pmm2 subunits (Supplementary Figure S4).

The structures of leisPMM showed within the same crystal, a combination of subunits in open (subunit A) and closed conformations (subunit B/C) (Kedzierski et al., 2006) (Supplementary Figure S6B). Interestingly, the leisPMM open conformation is similar to the closest conformation observed in Pmm2 subunit D_1_ (Supplementary Figure S6B). This comparison suggests that from the position observed in subunit D_1_, the core domain could rotate further to reach a similar closed position as seen in leisPMM (Figure 3C). If so, the subunit could switch from an open conformation as observed in the E1 subunit of Pmm2 to a closed conformation as in leisPMM, through a 30° rotation of the core domain, shortening the distance “d” between both domain from 22 Å to 9.5 Å (Figure 3C and Supplementary Figure S6B).

### Binding of the activator Gluc-1,6-P_2_

To obtain a more closed conformation of Pmm2, we co-crystallized the protein with the activator Glc-1,6-P_2_. The crystals diffracted X-rays to 2.75 Å resolution and were isomorphous with the type 1 apo crystal, containing six protein subunits organized as three independent dimers within the asymmetric unit (Table 1). In four subunits (B, C, D, and F), additional electron density was interpreted as the binding of Glc-1,6-P_2_ to the cap domain (Supplementary Figure S7A). However, the maps for the activator were less well-defined compared to the protein. Despite occupancy refinement reaching 73– 83%, the average temperature factors (B-values) for Glc-1,6-P_2_ were higher than those of the surrounding residues (Supplementary Figure S7B). These observations suggested that Glc-1,6-P_2_ is not tightly bound to the protein in the crystal.

The partial binding of Glc-1,6-P_2_ did not induce changes in the subunits, which showed conformations virtually identical to those observed in the type 1 crystal without activator (rmsd = 0.23– 0.75 for all Cα atoms), except subunit F that showed a small 6° rotation of the core domain (Supplementary Figure S8). Interestingly, the subunits displaying the most open conformations (A and E) did not seem to have the activator bound, whereas the electron density for the Glc-1,6-P_2_ molecule was more clearly defined in subunit D showing the closest conformation (Figure 3B and Supplementary Figure S7). This observation is reminiscent of the hPMM2 dimer structure, where Glc-1,6-P_2_ was bound to one subunit of the dimer and not to the other that exhibited a more open conformation (Briso-Montiano et al., 2022). Therefore, even though in both mouse and human PMM2 structures, Glc-1,6-bP_2_ interacts exclusively with the cap domain, the proximity of the core domain appears somehow important for its binding.

The binding of Glc-1,6-P_2_ closely resembles that observed in hPMM2 (Figure 4A,B). The activator lays against the hairpin β8-β9 in the cap domain, with the C1-phosphate binding to the nitrogen atom of Q173 and the side chains of S175, R137, and R130. Additionally, the hydroxyl groups of glucose are within hydrogen-bond distance of the side chains of residues R130, N124, R119, and D177. Conversely, the C6-phosphate extends toward the core domain but does not interact with the protein.

**Figure 4.**
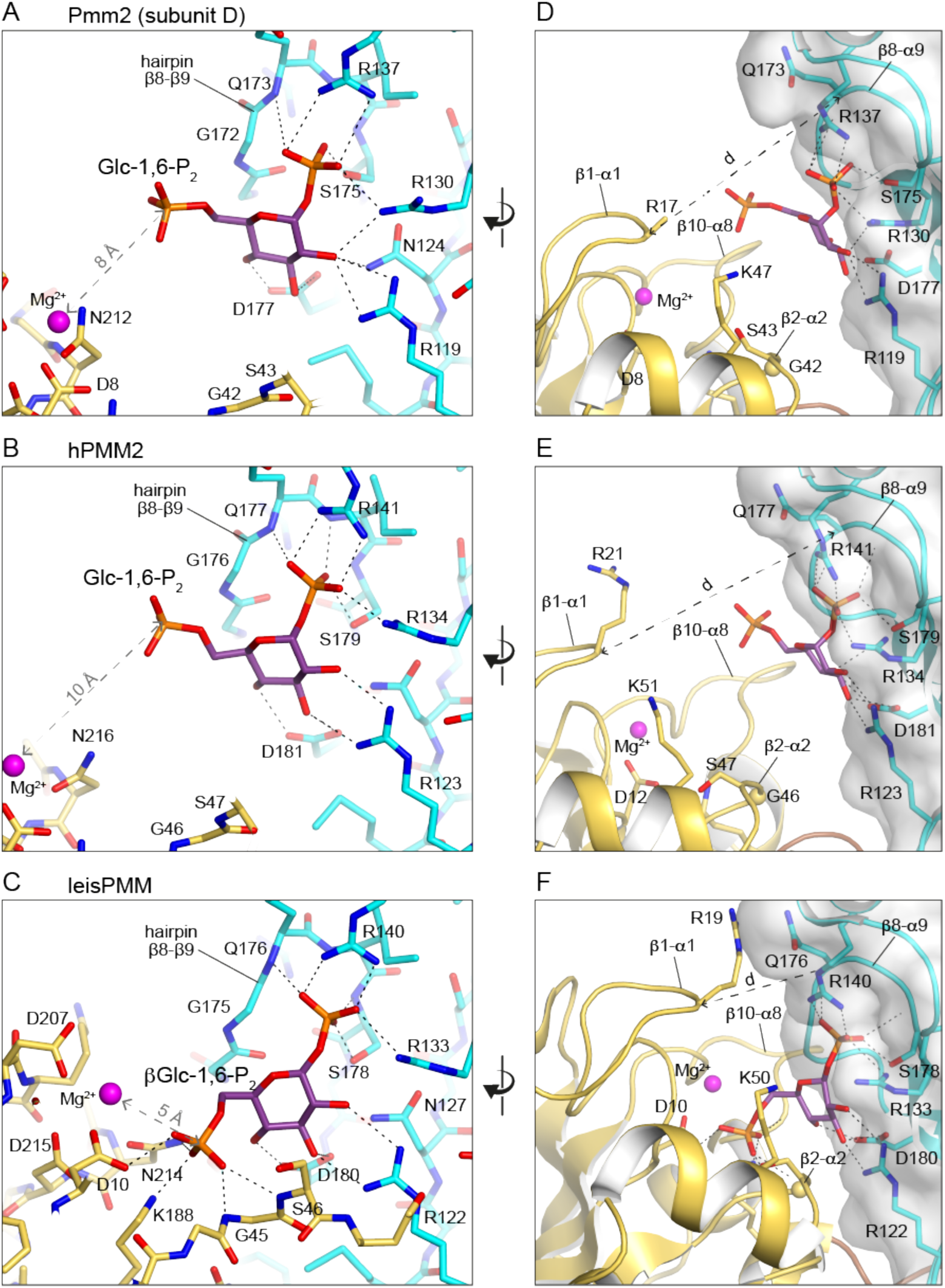
Binding of the activator Glc-1,6-P_2_. **A–C)** Detailed view of the binding of Glc-1,6-P_2_ to Pmm2-D_1_ **(A)** and hPMM2 **(B),** and of leisPMM in complex with the inert activator analog βGlc-1,6-P_2_ **(C)**. The ligand and interacting residues are shown in sticks, and the Mg-1 cation is represented in magenta spheres. Dashed black lines indicate H-bonds and ion pairs between the protein and the ligands. The protein carbon atoms are colored yellow and cyan for the core and cap domains, respectively. **D–F)** Rotated view showing the approximation of the core domain (in yellow) towards the ligand bound at the cap domain (in cyan). The dashed grey line indicates the distance between the Cα atoms of residues R21 (core domain) and Q176 (cap domain), also shown in Figure 3B.

This binding mode is similar to that seen in the crystal structure of leisPMM with β-glucose-1,6-bisphosphate (βGlc-1,6-P_2_), an inactive analog of the natural activator (Kedzierski et al., 2006; Monticelli et al., 2019). In leisPMM, the closure of the core domain facilitates additional interactions between the C6-phosphate and the nitrogen atoms of G45 and S46, as well as with the side chains of K188, D10, and N214 (Figure 4C). These residues in the core domain are conserved both in mouse and human PMM2, although they are out of reach of the activator in the more open conformation of the subunits (Figure 4D–F). Interestingly, an additional Cl^-^ anion near Mg-1 in the Pmm2 apo structures (Figure 2F) occupies a similar position in the core as the C6-phosphate in leisPMM, suggesting an affinity for negative charges that the C6-phosphate would fill upon core domain closure in PMM2. This subunit closure, along with the phosphate-binding –at the Cl^-^ site– via conserved active site elements, is likely responsible for the high-affinity binding of the activator, previously estimated for hPMM2 to have a K_D_ of 0.98 μM (Briso-Montiano et al., 2022).

### Molecular dynamic simulations with PMM2 dimer

To further explore the conformational flexibility of PMM2, we performed molecular dynamics (MD) simulations using the dimers of both mouse and human PMM2 in their apo states with the Cl^-^, Mg-1, and Mg-2 ions. In the initial models, both subunits within the dimer exhibited different conformations, none of which corresponded to the most open or closed positions observed in the crystal structures.

The MD simulations, conducted over 3 µs, showed a stable and rigid cap-cap dimer and extensive swinging motions of the projecting core domains (Figure 5A,B). Analysis of the rmsd values, comparing the structures generated along the MD trajectory with the initial model, indicated that both the core and cap domains maintained their compact globular structure throughout the simulation, but their relative orientation displayed a considerably wider range of conformational flexibility compared to the crystal structures (Figure 5C,D). The root-mean-square fluctuation (rmsf), which indicates the local positional variation for each residue, was considerably much lower for the cap domain than the core domain (Figure 5E). This underscores the view of the enzyme as a rigid cap-cap platform from which the core domains swing with a high degree of freedom.

**Figure 5.**
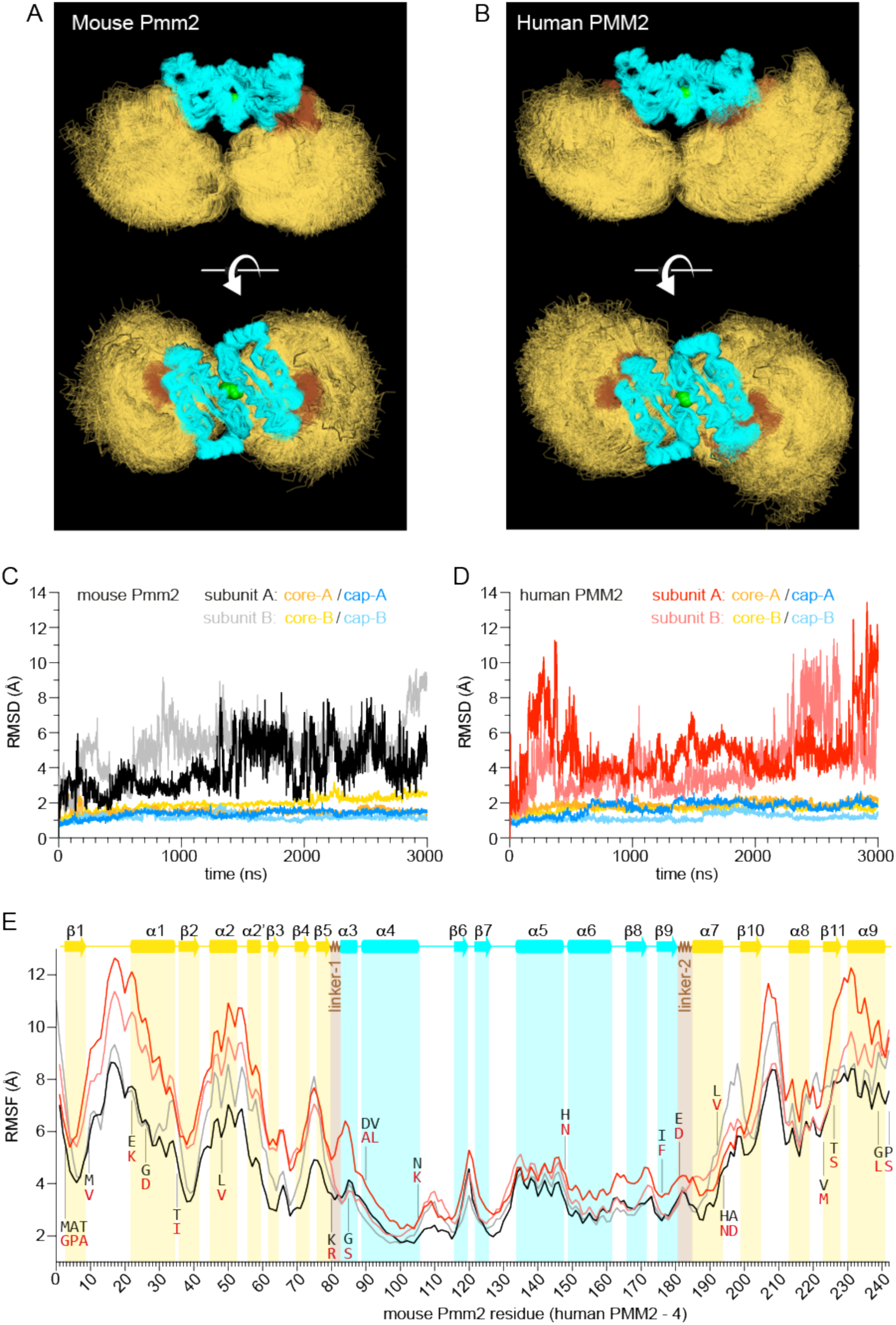
PMM2 molecular dynamic simulations. **A,B)** Perpendicular views of the superposition of 3,000 models extracted along the MD trajectory calculated for mouse and human PMM2. The cap and core domains are colored cyan and yellow, respectively, and linker regions are depicted in brown. The Cl^-^ anion is represented as a green sphere. Mg-1 and Mg-2 are not shown. **C,D)** RMSD for mouse and human PMM2 as obtained by the corresponding MD simulations. RMSD were calculated separately for each subunit in the dimer and for their respective cap and core domains. **E)** RMSF for the backbone atoms of the mouse (black and grey) and human (red and salmon) PMM2 subunits as obtained by the corresponding MD simulations. The x-axis indicates the residue number (n) for mouse PMM2, which in human PMM2 corresponds to the n-4 residue. The corresponding secondary structural elements are depicted above and beneath the graph, colored cyan and red for the cap and core domains, respectively. Positions that are not conserved among the two proteins are indicated.

The intersubunit Cl^-^ anion remained stably bound to both mouse and human enzymes during simulation, reinforcing its role as a key co-factor for PMM2 dimerization (Figure 5A,B). Similarly, Mg-1 remained firmly anchored at the active site, highlighting its essential role. In contrast, Mg-2 detached from Pmm2 subunits after 140 ns (subunit B) and 269 ns (subunit A) and from hPMM2 after 838 ns (subunit B) and 1012 ns (subunit A). Despite Mg-2 dissociation, the simulations showed no significant alterations in the core structure or increased domain flexibility. Indeed, hPMM2, which retained Mg-2 longer, showed greater overall flexibility. These findings prompted us to repeat the SEC experiments with 10 mM MgCl_2_ in the running buffer as mentioned earlier (Figure 2B). Under these conditions, the elution differences –attributed to different protein conformations– disappeared. Overall, these results suggest that Mg-2 plays a stabilizing function, and its loss may partially unfold the core domain, explaining the apparent larger protein size observed in SEC, though this effect is not seen within the MD simulation timescale.

The MD simulations revealed greater conformational flexibility than was inferred from the crystal structures. In the simulated trajectories, the hPMM2 dimer showed higher flexibility than Pmm2, with the core domains adopting more distant positions (Figure 5A,B). We measured the angle formed by the Cl^-^ anion and the catalytic Mg-1 ions in each subunit to track the position of the core domains relative to the cap-cap dimer (Figure 6A). Along the trajectory, the hPMM2 angles displayed a bimodal distribution, with a major peak at ∼85°, corresponding to a compact dimer with contacts between the core domains that involved polar interactions between helix α2’ (residues N58, D59, and E62) and loop α1-α2 (residues R21, Q22, and K26) (Figure 6B,D and Supplementary Figure S9). The second peak, centered at ∼125°, is a dimer with open subunits and no contact through the core domains, but different from that observed in the hPMM2 crystal structure, with an angle of 111°, which rarely appeared in the trajectory.

**Figure 6.**
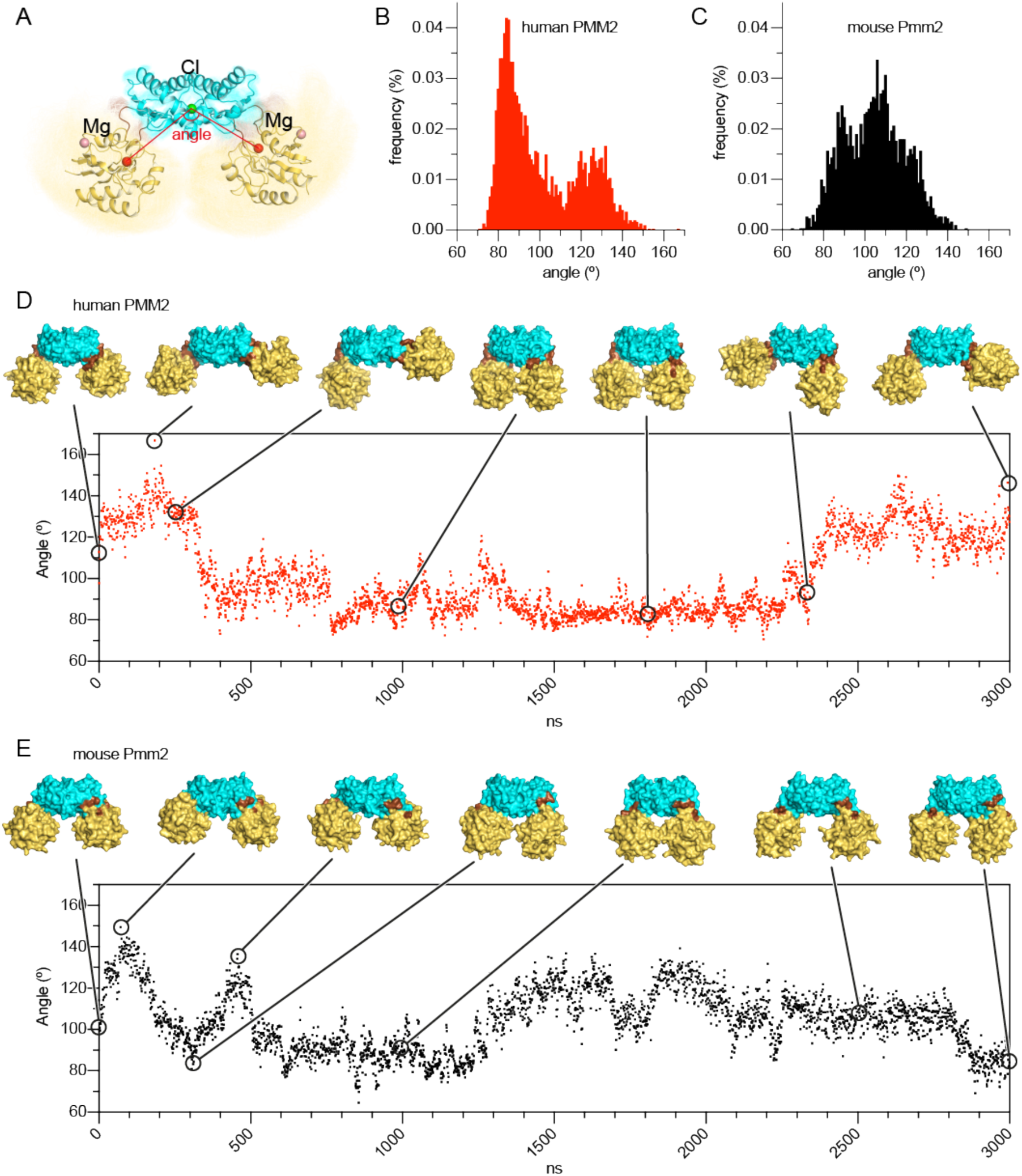
High conformational flexibility in the PMM2 dimer. **A)** Indication of the angle formed by the Cl^-^ anion in the dimerization interface and the Mg-1 cations in the active sites of both subunits. **B,C)** Distribution of the angle values as obtained by the MD simulations of human (B, in red) and mouse (C, black) PMM2 dimers. **D,E)** Value of the angles along the trajectory, showing representative models of the human (D) and mouse (E) PMM2 dimers.

On the other hand, for this particular simulation, Pmm2 fluctuated between a compact state (∼85°) and a conformation similar to the crystal structure (110°) without adopting the wide-open state observed in hPMM2 (Figure 6C,E). This difference likely reflects stochastic variability in the simulations, as domain and linker contacts are similar between both proteins.

Interestingly, despite the high flexibility of the core domain, the subunits did not adopt the closed conformation observed in leisPMM during the simulations, likely due to repulsion between positively charged side chains in the cap and core domains in the absence of the substrate or activator.

## Discussion

Enzymes are dynamic entities, and their movements are essential for their catalytic function. In this regard, our understanding of PMM2 remains limited, as we lack information about its reaction dynamics. Despite its apparent simplicity –a relatively small subunit with a bipartite active site– PMM2 catalyzes a complex reaction. The enzyme must accommodate three substrates –Glc-1,6-P_2_, Man6P, and Man-1,6-P_2_– at different stages of the reaction (Figure 1). This requires the cap domain to adjust for recognizing either the C1- or C6-phosphate and differently oriented hydroxyl groups, while ensuring that the opposite ends –C6- or C1-phosphates, or the C1-hydroxyl group– are correctly placed in the core domain for phosphoryl transfer. A greater challenge lies in the release and re-capture of the Man-1,6-P_2_ intermediate in a flipped orientation. Given is negligible concentration relative to the activator or substrate, the enzyme likely employs a mechanism to prevent complete dissociation into the solvent. Thus, efficient catalysis by PMM2 depends on a series of conformational changes to facilitate the orderly binding, release and reorientation of substrates.

Until now, the attempts to study PMM2 movements have been hindered by the constraints of crystallography. Both hPMM2 and its homolog hPMM1 crystallized exclusively in open conformations, even when bound to the activator or ligands (Briso-Montiano et al., 2022; Ji et al., 2018; Silvaggi et al., 2006). Similarly, leisPMM crystals displayed a mix of open and closed subunits, which remained unchanged upon binding an activator analog (Kedzierski et al., 2006). These crystal structures offered high-resolution yet static views apparently favored by lattice contacts and failed to capture ligand-induced movements or intrinsic protein flexibility.

In the present study, we aimed to expand our understanding of PMM2 catalytic motions by determining the structure of Pmm2, as its apparent size in solution suggested that it could crystallize in a catalytic competent closed conformation. The structures provide a higher-resolution model of the enzyme and its cofactors, along with eight distinct structural conformations, none of which corresponded to the anticipated closed state, as the protein crystallized in open conformations regardless of the binding of the activator. These results, combined with the comparison with other PMM structures, allowed us to define the enzyme’s architecture as a rigid, flat platform formed by the dimerization of the cap domains, with highly flexible core domains extending from each subunit. This structure depends on three ionic cofactors, occupying conserved positions across PMM structures: a Cl^-^ anion embedded in the cap-cap interface, essential for dimerization; a Mg^2+^ cation (Mg-1) firmly anchored at the phosphoryl transfer site in the core domain with a yet-to-be-defined catalytic function; and a second Mg^2+^ (Mg-2), whose structural role is corroborated in this work. Unlike the tightly bound Cl^-^ and Mg-1, Mg-2 is less stable and dissociates from the human and murine PMM2, as seen in MD simulations. While the MD timescale did not reveal the effects of Mg-2 dissociation, its conserved position and coordination suggest that it stabilizes the core domain. Indeed, the loss of Mg-2 likely leads to partial unfolding of the core domain, explaining the altered elution profiles of hPMM2 and Pmm2 in the SEC assays conducted with and without Mg^2+^ in the running buffer (Figure 2A).

The Pmm2 structures corroborate that the substrate binds to the cap domain via interactions with its phosphate and sugar ring, while its other end remains exposed, awaiting the closure of the core domain for proper alignment at the phosphoryl transfer site (Figure 4). Moreover, the superposition of the eight Pmm2 conformations suggests a putative directional movement of the core domain, potentially leading to the fully closed state seen in leisPMM crystals (Figure 3B,C). However, it remains unclear whether a simple “Pac-Man”-like opening and closing motion of the subunit is sufficient for the efficient progression through the reaction steps or if additional movements are required.

To better understand these mechanistic aspects and flexibility beyond the constraints of the crystal lattice, we performed MD simulations on the human and murine enzymes at an atomic scale. These simulations confirmed the enzyme’s architecture as a rigid cap-cap platform with highly flexible core domains. Surprisingly, the observed flexibility exceeded that inferred from crystal structures and appeared excessive for substrate exchange and reorientation, leaving its functional significance uncertain. This unexpected range of motion suggests a potentially more complex mechanism, possibly involving intersubunit communication within the dimer.

PMM2 is known to function exclusively as a dimer, with its most common pathogenic mutation, F119L (equivalent to F115 in Pmm2) –the residue coordinating the Cl^-^ co-factor (Figure 2E)– impairing dimerization (Andreotti et al., 2015, 2013). While dimerization is attributed primarily to protein stability, its potential functional role remains uncertain. Notably, although cooperative mechanisms have not been described, the cap-cap dimer features substrate-binding sites separated by ∼20 Å, a relatively short distance given that the reaction intermediate spans ∼9 Å. The flat cap-cap platform offers no physical barriers that impede the channeling or reorienting of substrates between active sites, supporting the idea of intersubunit communication. This hypothesis, previously suggested based on the leisPMM crystal structure (Kedzierski et al., 2006), may be enabled by the extensive movements of the flexible core domains.

A key challenge of this model is that it implies a shared reaction mechanism between subunits, suggesting that an inactivating mutation in one subunit would exert a dominant negative effect. However, all identified pathogenic hPMM2 variants are recessive, which seems to contradict this hypothesis. Nevertheless, the unexpected flexibility observed may indicate a previously unrecognized mechanism, warranting further investigation to elucidate how PMM2’s catalytic process unfolds.

MD simulations also revealed an unexpected and recurring dimer conformation with core-core interactions (Figure 6D,E). While this state is likely inactive, as core-core interactions prevent the closure over the cap domain, it may represent a resting, more resilient form of the enzyme. Importantly, no known pathogenic hPMM2 variants occur at the core-core interface. This suggests that this compact state may either lack functional relevance or resist point mutations, unlike the cap interface where F119L causes dimer dissociation. Regardless of its in vivo significance, the compact dimer observed in MD simulations could serve as a valuable target for pharmacochaperone development. Conformational-specific compounds that stabilize this state may enhance the stability of pathogenic PMM2 folding variants, preventing degradation and preserving residual enzymatic activity. Previous pharmacochaperone discovery efforts identified “compound VIII”, which exhibits activity against several pathogenic hPMM2 variants (Segovia-Falquina et al., 2022; Yuste-Checa et al., 2017). However, its mechanism of action remains unclear. Attempts to determine its binding site by X-ray crystallography were unsuccessful, possibly due to the compound’s poor solubility limiting crystal diffusion or weak affinity for the open dimer conformation enforced by crystal packing.

The compact dimer state offers an alternative framework for future drug development. Techniques like small-angle X-ray scattering (SAXS) could help identify compounds that stabilize specific conformations, such as the compact dimer, provided these states yield sufficiently distinct scattering profiles (Brosey et al., 2023). Furthermore, using the compact dimer as the initial model instead of the crystallized open structure could improve the rational or AI-guided ligand design to stabilize PMM2. An advantage of targeting the observed core-core interface is that pharmacochaperones will not compete with the substrate/activator for binding to the active site, potentially preserving the residual activity of the pathogenic variant.

In summary, this study underscores the high flexibility of the PMM2 dimer as a key factor for understanding its catalytic mechanism, the impact of pathogenic variants, and the potential for designing therapeutic compounds that stabilize more compact and resilient dimeric states.

## Materials and Methods

### Protein production

The sequence encoding mouse Pmm2 (UniProt Q9Z2M7) was PCR amplified from a pReceiver-B01 plasmid using Phusion High-Fidelity DNA Polymerase (New England Biolabs) and the pair of specific primers: 5’-<u>AAGTTCTGTTTCAGGGCCCG</u>ATGGCCACTCTCTGTCTCTTCG-3’ and 5’-<u>ATGGTCTAGAAAGCTTTA</u>TCAAGGGAAGAGCCCCTCACAG-3’. The primers contained regions (underscored) homologous to the ends of the pOPIN-B plasmid (Oxford Protein Production Facility) (Berrow et al., 2007) linearized with restriction enzymes KpnI and HindIII (New England Biolabs). The amplified insert and the digested plasmid were gel purified and ligated by *in vivo* DNA assembly (IVA cloning) (García-Nafría et al., 2016; Watson and García-Nafría, 2019). For this, insert and vector were mixed in water in a 2:1 molar ratio (62.4 ng insert and 162 ng plasmid) in a final volume of 5 µl, incubated on ice for 10 min, and used to transform *E. coli* XLBlue cells. The assembled plasmid (pOPINB-Pmm2) encoding mouse Pmm2 with an N-terminal hexahistidine tag and a PreScission protease site was verified by Sanger sequencing. BL21(DE3)pLysS cells transformed with pOPINB-Pmm2 were grown for 16 h in a shaking incubator at 37 °C in an autoinduction medium (Studier, 2005) supplemented with 100 µg·ml^-1^ ampicillin and 34 µg·ml^-1^ chloramphenicol. Cells were harvested by centrifugation and stored at −80°C. The bacterial pellet from 0.5 L culture was resuspended in 40 ml of buffer A (20 mM Tris-HCl pH 8, 0.5 M NaCl, 10 mM imidazole, 2 mM β-mercaptoethanol) with 0.1 mg·ml^-1^ Pefabloc protease inhibitor (Merck) and disrupted by sonication. The lysate was clarified by centrifugation at 23,700 g for 15 min. The supernatant was filtered through a 0.45 µm pore filter and loaded onto a 5 ml HisTrap HP affinity column (Cytiva) equilibrated in buffer A and connected to an NGC liquid chromatography system (Bio-Rad). Following extensive washing with buffer A supplemented with 25 mM imidazole, the protein was eluted with buffer A with 250 mM imidazole. Fractions found by SDS-PAGE to contain largely pure Pmm2 were mixed with GST-tagged PreScission protease (1:20 in a mass ratio relative to Pmm2) and dialyzed for 3 h versus dialysis buffer (20 mM Tris-HCl pH 8, 0.5 M NaCl, 25 mM imidazole, 2 mM β-mercaptoethanol). The digested and dialyzed sample was loaded a second time onto a HisTrap column coupled to a GST-trap 5 ml column (Cytiva) equilibrated in dialysis buffer to retain the noncleaved protein, the His-tag, and the protease. The untagged Pmm2 was not retained in the columns and was concentrated to ∼6 mg·ml^-1^ using an Amicon centrifugal ultrafiltration device (Merck) and loaded onto a HiLoad 16/60 Superdex 200 column (Cytiva) equilibrated in SEC buffer [20 mM Hepes pH 7.5, 0.2 M NaCl and 0.2 mM TCEP (tris(2-carboxyethyl)phosphine)]. Pmm2 eluted in a single peak and was concentrated to 6 mg·ml^-1^ as before and used directly for crystallization, or flash-frozen in liquid nitrogen and stored at −80 °C. All purification steps were carried out at 4 °C except SEC which was done at room temperature. Protein concentration was determined by the Bradford assay (Bio-Rad). The human PMM2 protein used in SEC experiments was obtained as previously reported (Briso-Montiano et al., 2022).

### Crystallization

Initial crystallization screenings were performed at 18 °C in 96-well MRC crystallization plates (Molecular Dimensions) with drops of 1 μl protein at 5 mg·ml^-1^ and 1 µl reservoir solution and 60 μl of reservoir solution from the Index, JCSG+ and PACT commercial screenings (Qiagen). Initial hits were optimized in hanging drop 24-well plates (Hampton Research) mixing 2 µl of protein with or without 2 mM α-glucose-1,6-bisphosphate (Sigma) and 2 µl of reservoir solution. Best crystals appeared after 2-7 days in 0.1 M Tris-HCl pH 8.5, 0.2 M MgCl_2_, and either 20-22% PEG 8000 or 27-30% PEG 3350. Crystals were cryo-protected by briefly soaking in mother liquor with increasing concentrations of PEG up to 30% and 5% glycerol. Crystals were fished with cryo-loops and flash-cooled in liquid nitrogen.

### Data collection and structure determination

X-ray diffraction data were remotely collected at ESRF beamline ID30B (DOI 10.15151/ESRF-ES-649779879) using MXCuBE (Gabadinho et al., 2010) and at ALBA beamline BL13-XALOC. For each dataset, a total wedge of 180° was collected with 0.1-0.15° oscillation and approximately 0.1 s exposure per frame using Pilatus 6M detectors. Data processing and scaling were performed automatically with XDS (Kabsch, 2010) and autoPROC (Vonrhein et al., 2011). Crystallographic phases were obtained by molecular replacement using PHASER (McCoy et al., 2007) and human PMM2 (PDB ID 7O0C; (Briso-Montiano et al., 2022) as the search model. Final models were obtained by iterative cycles of Coot (Emsley et al., 2010) building and Phenix (Adams et al., 2010) or Refmac (Murshudov et al., 2011) refinement using CCP4 (Winn et al., 2011). Structural flexibility and subunit movements were analyzed with CCP4 and DynDom (Veevers and Hayward, 2019). Structures were represented with PyMOL (Schrödinger; https://pymol.org/2/).

### Molecular dynamics

MD simulations of PMM2 dimers were performed using as starting models, the X-ray structures of mouse PMM2 from the type 2 crystal and of the human enzyme (PDB ID 7O0C) without bound Glc-1,6-P_2_. MD simulations were performed with the GROMACS 2022.6 software package (Abraham et al., 2015). Initial models and files for GROMACS were prepared with CHARMM-GUI (www.charmm-gui.org) (Jo et al., 2008). For MD simulations, the CHARMM36m force field was used (Huang et al., 2017). The protein was solvated with water and counterions (150 mM KCl) were added by replacing a corresponding number of water molecules to achieve a neutral condition. Van der Waals interactions were smoothly switched off at 10–12 Å by a force-switching function and long-range electrostatic interactions were treated with the particle mesh Ewald method (Darden et al., 1997). The simulations were performed at 310.15 K using a Nose-Hoover thermostat with τT = 1.0 ps. Pressure was maintained constant at 1 bar with a Parrinello-Rahman algorithm with a semiisotropic coupling constant τP = 5.0 ps and compressibility = 4.5 Å∼ 10^−5^ bar^−1^. The LINCS method was used to constrain bond lengths. A time step of 2 fs was used for numerical integration. Coordinates were saved every 5 ps for analysis and the total simulation time was 3,000 ns. Analysis was performed using the GROMACS suite programs. The trajectories and generated models were visualized with PyMol.

## Acknowledgments

We acknowledge the European Synchrotron Radiation Facility (Grenoble, France) and ALBA synchrotron (Barcelona, Spain) for the provision of beam time on ID30B and BL13-XALOC, and we would like to thank Gianluca Santoni (ESRF) and Xavier Carpena and Isidro Crespo (ALBA) for assistance. This work was supported by the following grants: RTI2018-098084-B-100 financed by the Spanish Ministerio de Ciencia e Innovación/Fondo Europeo de Desarrollo Regional (AEI/FEDER UE) to SR-M; grant PID2021-128468NB-I00 financed by MCIN/AEI/10.13039/501100011033 to SR-M; grant PI19/01155 to BP; Fundación Ramón Areces Ciencias de la Vida, XX National call, to SR-M; CIBER of Rare Diseases (CIBERER) Innovative Therapies Award Number ERT18TRL746, to BP; Regional Government of Madrid grant CAM, B2017/BMD3721 to BP; and by granted access to ALBA (proposals 2021075216) and ESRF (proposal MX-2351) synchrotrons to SR-M. During the development of this study, FdC-O was a postdoctoral fellow of the Regional Government of Valencia (APOSTD2021/099 Generalitat Valenciana).

## Notes

### Competing Interest Statement

The authors have declared no competing interest.

