## Supplementary Figures for "High conformational flexibility of phosphomannomutase 2: Implications for functioning mechanisms, stability and pharmacological chaperone design"

---

*Corresponding author

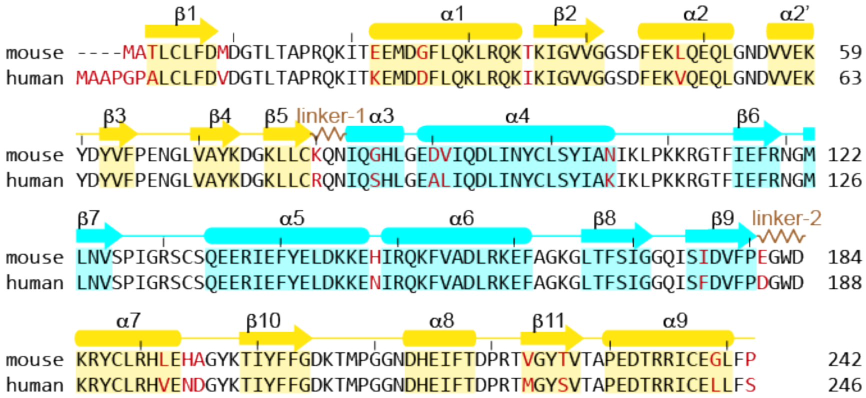


**Supplementary Figure S1. Human and mouse PMM2 alignment.** Sequence alignment of mouse and human PMM2 with secondary structural elements indicated on top. Non-identical residues are shown in red.

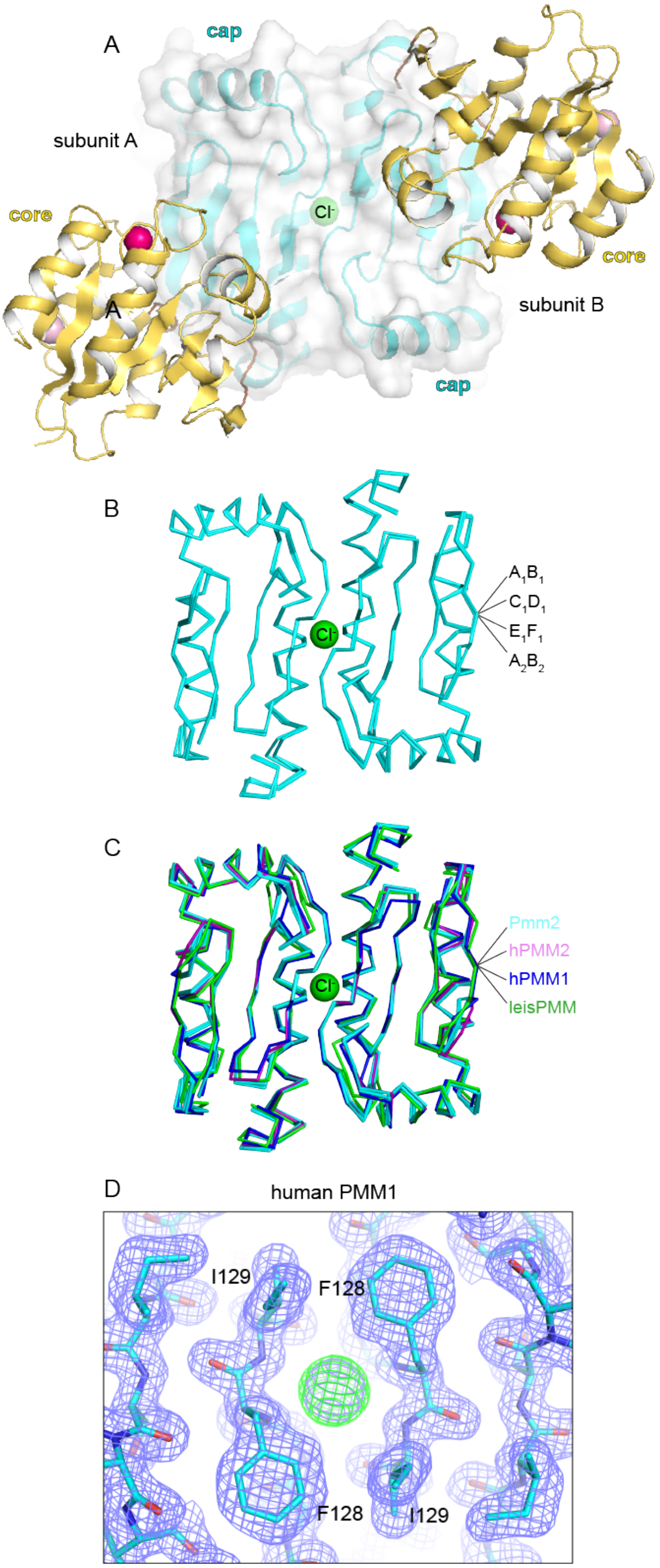


**Supplementary Figure S2. The cap-cap dimer is a rigid platform with an embedded $Cl^-$ anion. A)** Representation of the Pmm2 dimer in a perpendicular view to that shown in Figure 2D. **B)** Ribbon representation of the superposed cap domains in the four Pmm2 cap-cap dimers contained in the asymmetric units of type 1 and type 2 crystals. The $Cl^-$ anion is represented as a green sphere. **C)** Superposition of the cap-cap dimers from Pmm2, hPMM2, hPMM1, and leisPMM. **D)** Detail of the crystal structure of hPMM1 (PDB ID 2FUE) with the $2F_{obs}$-$F_{calc}$ electron density map contoured at 1.0 $\sigma$ and shown as a blue mesh. The $F_{obs}$-$F_{calc}$ map contoured at 2.5 $\sigma$ is colored green and indicates the conserved position of the $Cl^-$ anion that was not included in the final model.

| | | crystal type 1 | | | | | | crystal type 2 | |
|---|---|---|---|---|---|---|---|---|---|
| | | $A_1$ | $B_1$ | $C_1$ | $D_1$ | $E_1$ | $F_1$ | $A_2$ | $B_2$ |
| crystal type 1 | $A_1$ | | 0.801 | 1.929 | 2.106 | 1.187 | 1.462 | 0.719 | 2.023 |
| | $B_1$ | | | 1.451 | 1.619 | 1.386 | 1.068 | 0.600 | 1.600 |
| | $C_1$ | | | | 0.587 | 2.403 | 0.832 | 1.454 | 1.460 |
| | $D_1$ | | | | | 2.656 | 1.065 | 1.645 | 1.278 |
| | $E_1$ | | | | | | 1.793 | 1.304 | 2.888 |
| | $F_1$ | | | | | | | 1.048 | 1.747 |
| crystal type 2 | $A_2$ | | | | | | | | 1.782 |
| | $B_2$ | | | | | | | | |

**Supplementary Figure S3. Pmm2 superpositions.** RMSD values for superposition among the eight Pmm2 subunits observed in the type 1 and 2 crystals. Larger differences are colored in darker red.

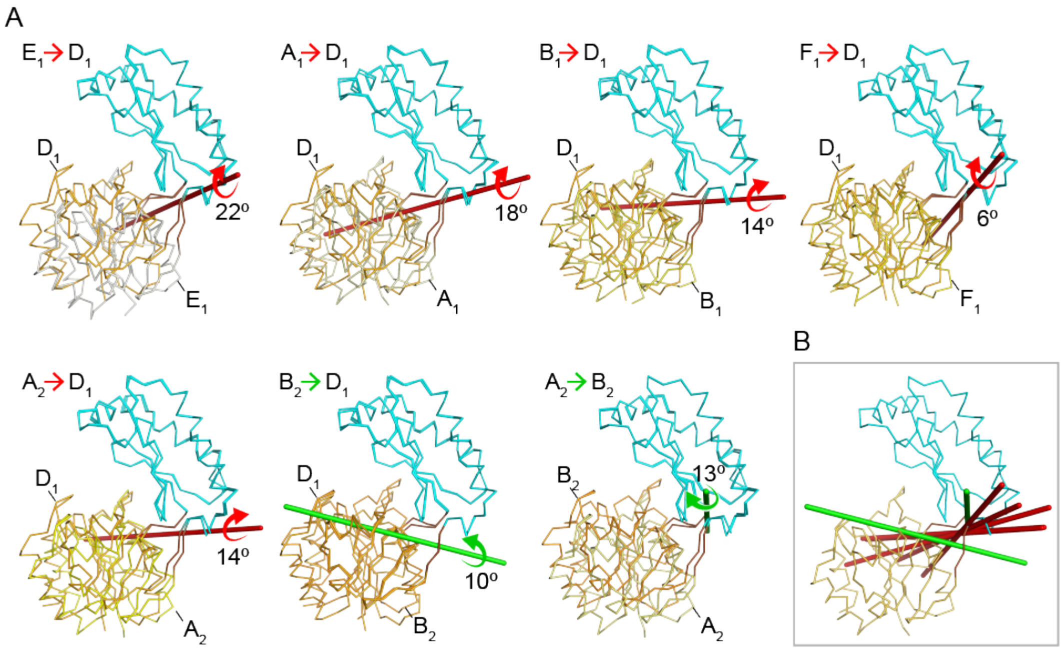


**Supplementary Figure S4. Pmm2 conformational movements within the crystals. A)** One-to-one superposition through the cap domain of the different Pmm2 subunits. The red or green rods indicate the closure axis for the rotation needed to superpose the core domains, which is indicated with a curved arrow. **B)** Superposition of the different closure axes shown in A.

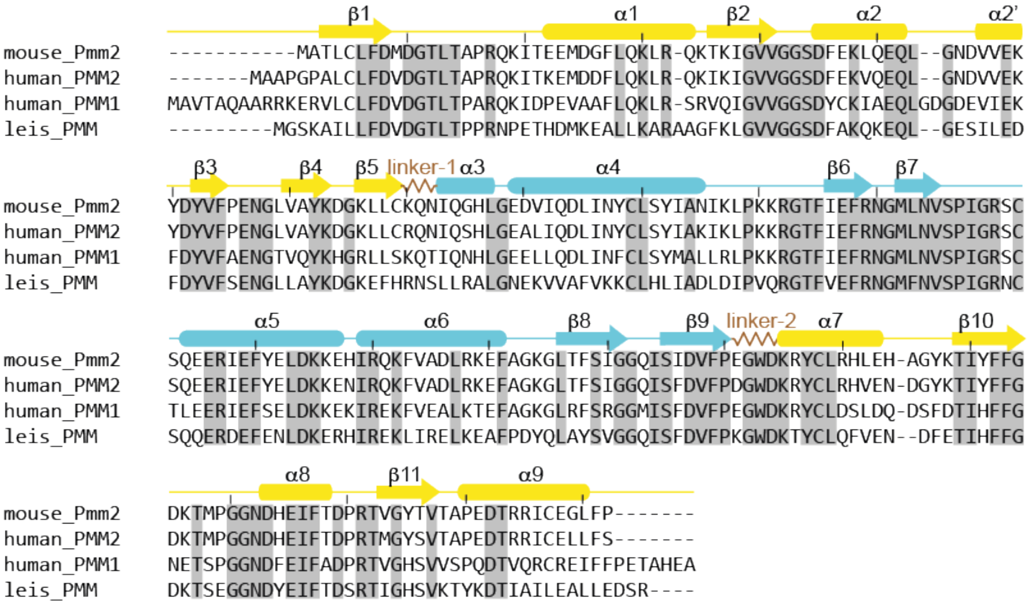

**Supplementary Figure S5. Sequence conservation among PMMs.** Sequence alignment of Pmm2 (UniProt A9Z2M7), hPMM2 (O15305), hPMM1 (Q92871) and leisPMM (Q95ZD7), for which crystal structures are available. Secondary structure elements are depicted above the sequences and colored yellow and cyan for the core and cap domains, respectively. Identical residues are shown with a grey background.

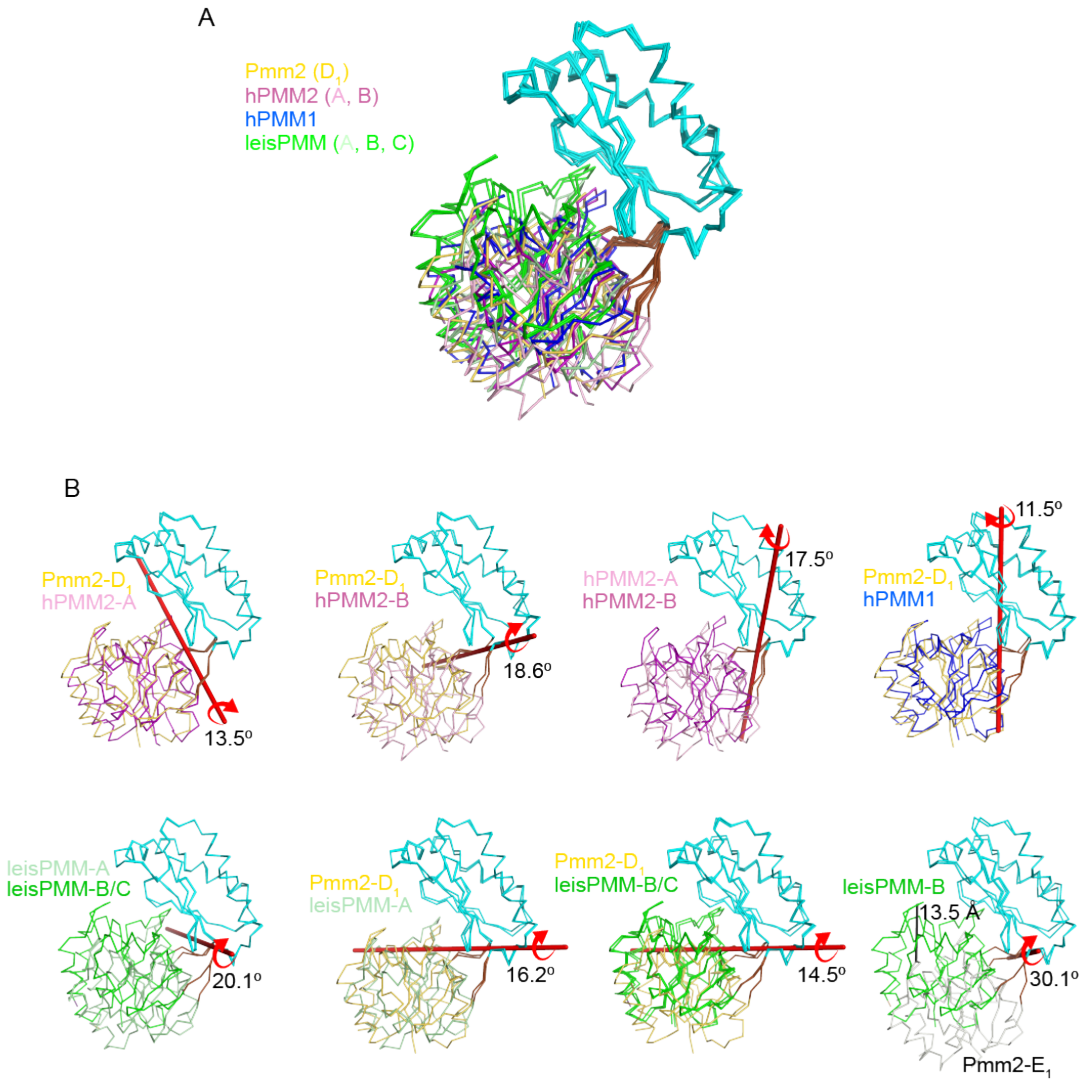


**Supplementary Figure S6. Structural comparison between Pmm2, hPMM2, hPMM1 and leisPMM. A)** Ribbon representation of the superposition of Pmm2 subunit $D_1$, hPMM2 (subunits A and B; PDB entry 7O58), hPMM1 (PDB entry 2FUE), and leisPMM (subunits A, B, and C; PDB 2I55) through the cap domains (colored in cyan), showing the different orientations of the core domains, which are depicted in different colors. **B)** Superposition through the cap domains of the indicated protein subunits with Pmm2-$D_1$. The subunits are colored as in **A**. The thick rods indicate the rotation axes for the superposition of the core domains.

A

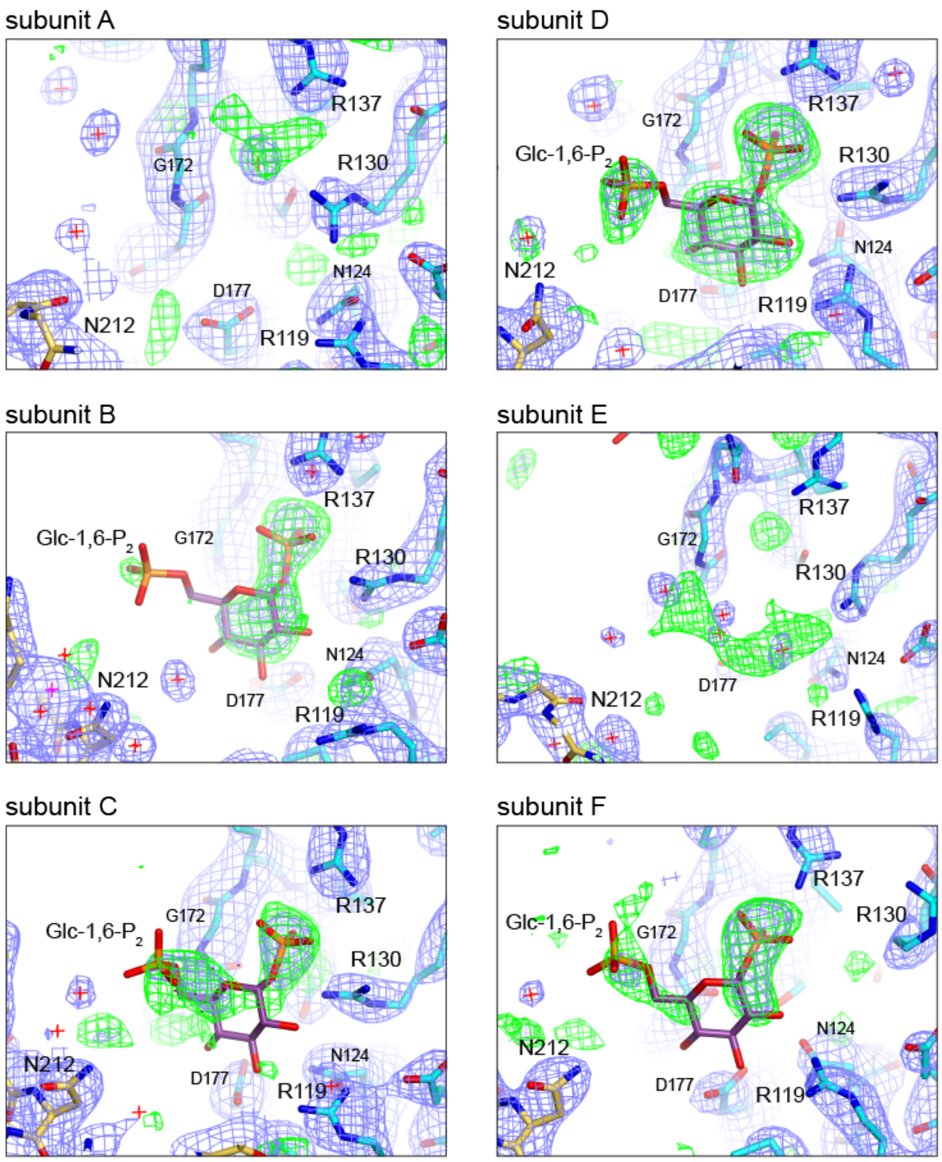


B

| subunit | Glc-1,6-$P_2$ occupancy | average B-factor | | | | | | |
|---|---|---|---|---|---|---|---|---|
| | | Glc-1,6-$P_2$ | R119 | R130 | R137 | G172 | Q173 | protein |
| A | - | - | 59 | 58 | 62 | 49 | 57 | 50 |
| B | 73 % | 83 | 64 | 61 | 61 | 61 | 61 | 60 |
| C | 80 % | 99 | 64 | 68 | 74 | 78 | 79 | 58 |
| D | 83 % | 77 | 56 | 58 | 58 | 58 | 55 | 65 |
| E | - | - | 80 | 85 | 105 | 88 | 94 | 78 |
| F | 81 % | 109 | 79 | 98 | 106 | 99 | 101 | 82 |

**Supplementary Figure S7. Binding of the activator Glc-1,6-$P_2$ in Pmm2 type 1 crystal. A)** Detailed view of the six Pmm2 subunits in crystal type 1 grown in the presence of 2 mM Glc-1,6-$P_2$. The $2F_{obs}$-$F_{calc}$ map (in blue) and the $F_{obs}$-$F_{calc}$ map (in green) are contoured at 1.0 $\sigma$ and 2.5 $\sigma$, respectively, and were calculated by excluding the activator from the protein model. **B)** Refined occupancy of the Glc-1,6-$P_2$ bound to the different subunits, and average temperature factors (B-factor) for the activator and the interacting residues, and for the entire protein subunit.

A

| subunit | G16bP | rmsd (Å) | C$\alpha$'s |
|---|---|---|---|
| A | – | 0.23 | 241 |
| B | + | 0.35 | 241 |
| C | + | 0.28 | 241 |
| D | + | 0.58 | 241 |
| E | – | 0.42 | 241 |
| F | + | 0.75 | 241 |

B

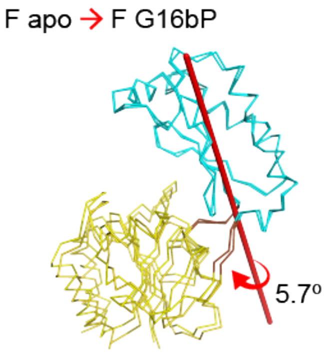


**Supplementary Figure S8. Binding of the activator does not induce conformational changes. A)** RMSD for the superposition of the Pmm2 subunits crystallized (type 1 crystal) with and without Glc-1,6-$P_2$. The higher rmsd values for the superposition of subunit F is highlighted in red. **B)** Ribbon representation of the superposition through the cap domains of the F subunits with and without Glc-1,6-$P_2$. The red rod and arrow indicate the ~6º rotation of the core domain between both structures.

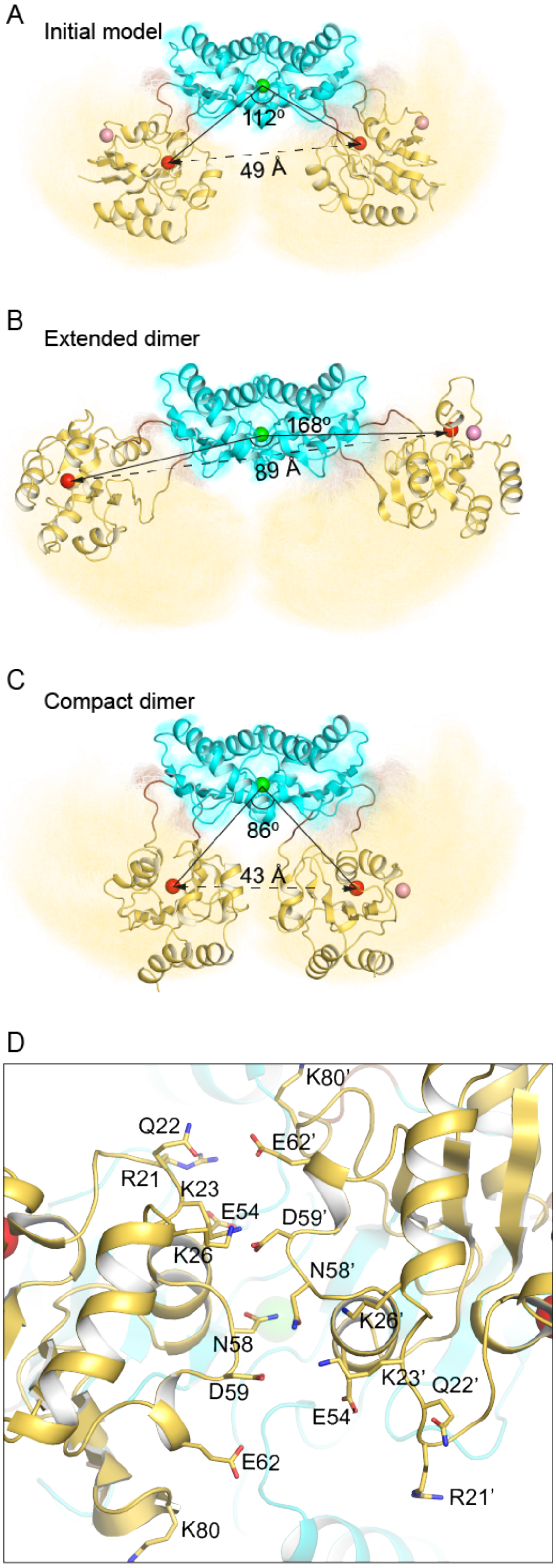


**Supplementary Figure S9. Flexibility of the PMM2 dimer and core-core domain interactions. A–C)** Cartoon representation of hPMM2 dimers along the MD trajectories: A) initial stage, as observed in the crystals; (B) extended conformation; and (C) compact state. The angle between the $Cl^-$ and Mg-1 ions and the distance between the Mg-1 ions are indicated. The structures are represented above a ribbon representation of the superposed snapshots of the model extracted every ns along the 3.000 ns trajectory. **D)** Detailed view of the interactions between the core domains in one of the compact dimer conformations observed in the MD trajectory.
